# Graph Neural Networks Model Based on Atomic Hybridization for Predicting Drug Targets

**DOI:** 10.1101/2025.11.19.689219

**Authors:** Ashraf Mohamed, Noha Galal, Bernard R. Brooks, Muhamed Amin

**Affiliations:** AI Department, Areeb Innovative Technologies, Heliopolis, Cairo 11811, Egypt; Centre for Theoretical Physics, The British University in Egypt, P.O. Box 43, El Sherouk City, Cairo 11837, Egypt; Biomedical Sciences Program, University of Science and Technology, Zewail City of Science and Technology, Giza, 12578, Egypt; Laboratory of Computational Biology, National Heart, Lung and Blood Institute, National Institutes of Health, Bethesda, Maryland 20892, USA; Department of Sciences, University College Groningen, University of Groningen, 9718 BG Groningen, The Netherlands

## Abstract

Accurate prediction of half-maximal inhibitory concentration (IC_50_) values is critical for accelerating drug discovery, yet traditional quantitative structure-activity relationship (QSAR) models often have limited ability to capture both local structural patterns and global physicochemical properties essential for bioactivity. We developed a hybrid deep learning framework that integrates graph neural networks with explicit molecular descriptors to address this limitation. The model learns from molecular graphs encoding atomic and bond features while incorporating interpretable physicochemical properties and structural fingerprints. Trained and validated on 14,316 compounds across nine diverse biological targets including kinases, nuclear receptors, and proteases, our approach achieved an overall test R^2^ of 0.87, consistently outperforming previously reported methods by 6-42% across evaluated targets. The model demonstrated robust generalization with near-identical training and test performance, while maintaining partial interpretability through transparent descriptor contributions and attention mechanisms. By synergistically combining data-driven learning with domain knowledge, this hybrid framework offers improved accuracy and interpretability for structure-activity modeling, facilitating more efficient compound prioritization and optimization in early-stage drug discovery programs.

## INTRODUCTION

Accurately predicting the bioactivity of small molecules, particularly their half-maximal inhibitory concentration (IC_50_), is a cornerstone of rational drug discovery and development. IC_50_ values provide a quantitative measure of a compound’s potency in inhibiting a biological process, making them essential for compound prioritization, hit-to-lead optimization, and toxicity assessment. Traditional quantitative structure–activity relationship (QSAR) models have long been used for this task, relying on predefined molecular descriptors or structural fingerprints derived from chemical intuition and empirical observations^1–5^. While these models have shown success in narrow chemical spaces, their sole dependence on handcrafted features often limits their generalizability and robustness when applied to diverse or novel chemical scaffolds^6–8^.

Recent advances in machine learning, and deep learning in particular, have led to a paradigm shift in cheminformatics. Among these, graph neural networks (GNNs) have gained prominence due to their ability to directly operate on molecular graph structures and learn task-specific features in an end-to-end fashion^9,10^. In GNN-based molecular modeling, atoms are typically represented as nodes and chemical bonds as edges, with associated node and edge features encoding atom types, hybridization states, formal charges, aromaticity, bond orders, and other chemically relevant properties^11^. This formulation enables GNNs to capture local atomic environments and topological features of molecules without explicit feature engineering.

Several studies have demonstrated the effectiveness of GNNs in molecular property prediction tasks. Gilmer et al.^12^ introduced the message-passing neural network (MPNN) framework, which unifies many GNN architectures and applies them to quantum chemistry problems, showing state-of-the-art performance on several benchmarks. Building on this, other work have extended GNNs to predict solubility^13^, toxicity^14^, protein–ligand binding affinity^15^, biological activity^16^ and other general purpose studies^17–22^. Notably, these models often outperform traditional descriptor-based QSAR models, particularly when trained on large, diverse datasets.

Despite these successes, conventional GNNs may still underutilize certain chemically meaningful concepts. One such example is atomic hybridization, which is typically treated as just another categorical feature appended to the node attribute vector. However, hybridization plays a crucial role in determining molecular geometry, orbital overlap, and reactivity–factors that directly impact molecular recognition and bioactivity^23–27^. Treating hybridization merely as a side feature may limit its influence on molecular function and limit the representational capacity of the model. To address this limitation, we propose a hybridization-explicit graph representation. Unlike standard approaches that treat atomic element and hybridization as separate features, our model utilizes a combinatorial node embedding strategy wherein node identities are defined by unique element-hybridization tuples. We define node identities not merely by atomic symbol (e.g., C, N, O), but by unique element-hybridization tuples (e.g., distinguishing C_SP_^3^ from C_sp_^2^ as distinct node classes). This creates a fine-grained topological representation where the electronic configuration and spatial orientation of atoms are encoded directly into the fundamental node vocabulary.

We integrate this enriched graph representation into a Hybrid GNN-Descriptor Architecture. In our approach, one branch utilizes a Graph Attention Network (GAT) to process these high-fidelity molecular graphs, while a parallel branch explicitly models key physicochemical descriptors (e.g., LogP, Molecular Weight). By integrating these two distinct modalities via a fusion layer, we aim to capture both the detailed electronic environments of individual atoms and the global physicochemical constraints of the molecule. We evaluate our approach using a curated dataset of small molecules with experimentally determined IC_50_ values, applying a dual-branch architecture where a graph attention network (GAT)-based encoder processes the hybridization-based graphs while a parallel network encodes established physicochemical descriptors. Our results demonstrate that this chemically informed representation yields improved predictive accuracy and stronger correlation with experimental IC_50_ values compared to traditional atom-based GNNs and fingerprint-based baselines. Furthermore, the model exhibits enhanced generalization, particularly on structurally diverse compounds, highlighting the benefit of incorporating domain knowledge into graph construction. We explicitly benchmark our Hybrid model against well-established baselines, including traditional QSAR and pure GNNs, using our own curated dataset. Furthermore, we provide a comprehensive architectural comparison with state-of-the-art literature models such as GraphTCDR^19^, XGDP^28^, and MTATFP ^29^. We note that while these advanced models often utilize different datasets or multi-modal inputs (e.g., cell-line omics), preventing a direct quantitative comparison, this architectural analysis is crucial for positioning our model’s unique feature integration and design philosophy for ligand-based virtual screening.This work contributes to the growing field of interpretable and domain-aware molecular machine learning. By integrating hybridization as a fundamental topological element, our method offers a novel perspective on molecular graph representation and underscores the value of chemically meaningful abstractions in deep learning pipelines. As the field continues to explore new GNN architectures and data modalities, such chemically enriched graph topologies may play a crucial role in improving prediction accuracy and mechanistic interpretability in drug discovery applications^30,31^.

## METHODS

### 1. Dataset Preparation

The data used in this study was retrieved from ChEMBL database^32,33^. We selected nine diverse biological targets spanning different protein families to assess model generalizability: CHEMBL203, CHEMBL204, CHEMBL206, CHEMBL1827, CHEMBL2835, CHEMBL2842, CHEMBL332, CHEMBL4302, and CHEMBL5763. For each target, we retrieved all available compound activity measurements with standard type IC_50_ using the ChEMBL web services API.

The raw data underwent systematic preprocessing to ensure quality and consistency. Molecular structures were represented as SMILES (Simplified Molecular Input Line Entry System) strings, which provide a compact text-based encoding of molecular structure. We validated all SMILES strings using RDKit^34^ to confirm they could be successfully parsed into valid molecular structures, removing any entries with malformed or ambiguous SMILES representations. IC_50_ values were extracted and filtered to retain only numeric measurements with appropriate units (typically nanomolar or micromolar concentrations). Entries with missing, non-numeric, or ambiguous IC_50_ values were excluded from further analysis.

Following initial filtering, we applied additional quality control steps to improve dataset reliability. IC_50_ values less than or equal to zero were removed, as these represent invalid measurements that cannot be logarithmically transformed. To handle duplicate measurements of the same compound against a given target, we performed deduplication based on canonical SMILES representations, retaining only unique compound-target pairs. When multiple IC_50_ measurements existed for the same compound, we retained the median value to reduce the impact of experimental variability.

All IC_50_ values were logarithmically transformed (log10 scale) to normalize their distribution, as bioactivity measurements typically follow a log-normal distribution spanning several orders of magnitude. This transformation also aligns with how potency data is reported. The resulting log(IC_50_) values served as the target variable for model training, with lower values indicating higher potency.

All nine ChEMBL target datasets were pooled together to create a combined multi-target training set, allowing the model to learn general structure-activity relationships that transfer across different biological targets. Dataset sizes after preprocessing ranged from approximately 500 to 8,000 compound-activity pairs per target. The combined dataset was randomly split into 80% training and 20% test sets using a fixed random seed (42) to ensure reproducibility. To address class imbalance in the training set, we employed oversampling of underrepresented targets to ensure balanced learning across all protein families. The test set was held out during all model development and hyperparameter tuning to provide an unbiased estimate of generalization performance, with performance evaluated both overall across all test samples and separately for each individual target.

### 2. Model Architecture

#### Hybrid Graph Neural Network with Molecular Descriptors

We developed a hybrid deep learning architecture that integrates graph neural networks (GNNs) with traditional molecular descriptors for IC_50_ prediction. The model consists of two parallel branches that capture complementary aspects of molecular structure and properties. A block diagram illustrating this architecture and the functional role of each component is provided in Figure 1.

**Figure 1.**
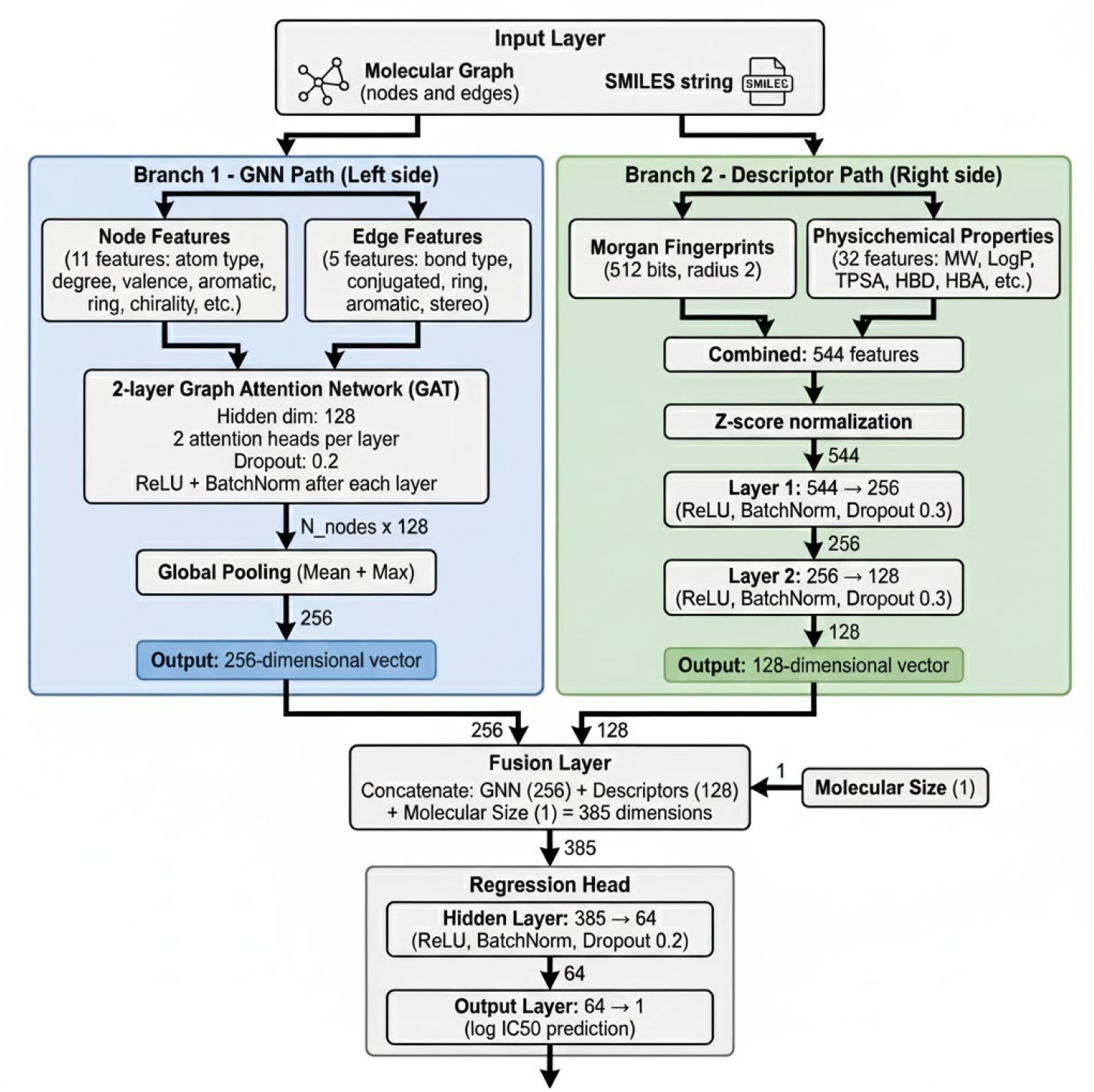
Architecture of a Hybrid Graph Neural Network with Molecular Descriptors for IC_50_ Prediction. The model integrates two parallel branches: a graph neural network branch and a molecular descriptor branch. The GNN branch processes molecular graphs with node features (X, 11 dimensions including atom type, hybridization, atomic number, degree, formal charge, hydrogen count, aromaticity, ring membership, chirality, ring size, and total valence) and edge features (5 dimensions including bond type, conjugation, ring membership, stereochemistry, and aromaticity). Node and edge features are embedded and passed through a two-layer Graph Attention Network (GAT) with 128 hidden dimensions and 2 attention heads per layer. The resulting node representations are aggregated via global mean and max pooling operations, producing a 256-dimensional graph embedding. In parallel, the descriptor branch computes 544 molecular features directly from SMILES strings, consisting of 512-bit Morgan fingerprints (radius 2) and 32 physicochemical properties (including lipophilicity, hydrogen bonding capacity, molecular structure, electronic properties, and shape descriptors). These descriptors are processed through a two-layer feedforward neural network (544 → 256 → 128 dimensions) with batch normalization and dropout (0.3). The 256-dimensional GNN embedding, 128-dimensional descriptor embedding, and a single molecular size feature (sum of atomic numbers) are concatenated to form a 385-dimensional fused representation. This combined feature vector is passed through a single-layer regressor (385 → 64 → 1) with batch normalization and dropout (0.2) to produce the final log(IC_50_) prediction.

#### Graph Neural Network Branch

The GNN branch processes molecular graphs where atoms are represented as nodes and bonds as edges. Each molecule is converted from SMILES notation to a molecular graph representation using RDKi^t34^. For each atom node, we encode eleven features capturing atomic identity, connectivity, and chemical environment. These include the combined atom type and hybridization state through categorical encoding, atomic number, atom degree representing the number of neighbors, and formal charge. To capture hydrogen bonding potential, we include the total hydrogen count combining both implicit and explicit hydrogens. Topological features include binary indicators for aromaticity and ring membership. Stereochemical information is preserved through binary flags for R and S chirality configurations. Additionally, we encode the size of the smallest ring containing each atom, with a value of zero for non-ring atoms. Finally, the total valence provides information about the bonding capacity of each atom.

For bond edges, we extract five features characterizing the connection between atoms. The bond type distinguishes between single, double, triple, and aromatic bonds. Binary features indicate whether the bond is conjugated, part of a ring system, or aromatic. Stereochemical configuration of the bond is also encoded to preserve spatial information relevant to biological activity.

The molecular graph is processed through a two-layer Graph Attention Network (GAT)^35^ with a hidden dimension of 128 and two attention heads per layer. The GAT layers employ attention mechanisms to learn the importance of neighboring atoms when updating node representations, allowing the model to focus on structurally and chemically relevant regions of the molecule. Dropout regularization of 0.2 is applied after each GAT layer to prevent overfitting, along with ReLU activation and batch normalization. Following graph convolution, we generate graph embeddings using both global mean and maximum pooling operations, producing a 256-dimensional vector that captures both the average molecular features and the most salient structural characteristics.

#### Molecular Descriptor Branch

The descriptor branch computes physicochemical and structural properties directly from SMILES strings also using RDKit^34^. We selected 32 interpretable molecular descriptors known to correlate with drug activity and bioavailability. These descriptors span five categories that capture complementary aspects of molecular structure and properties. Table 1 summarizes the full list of descriptors used in this study.

**Table 1.**
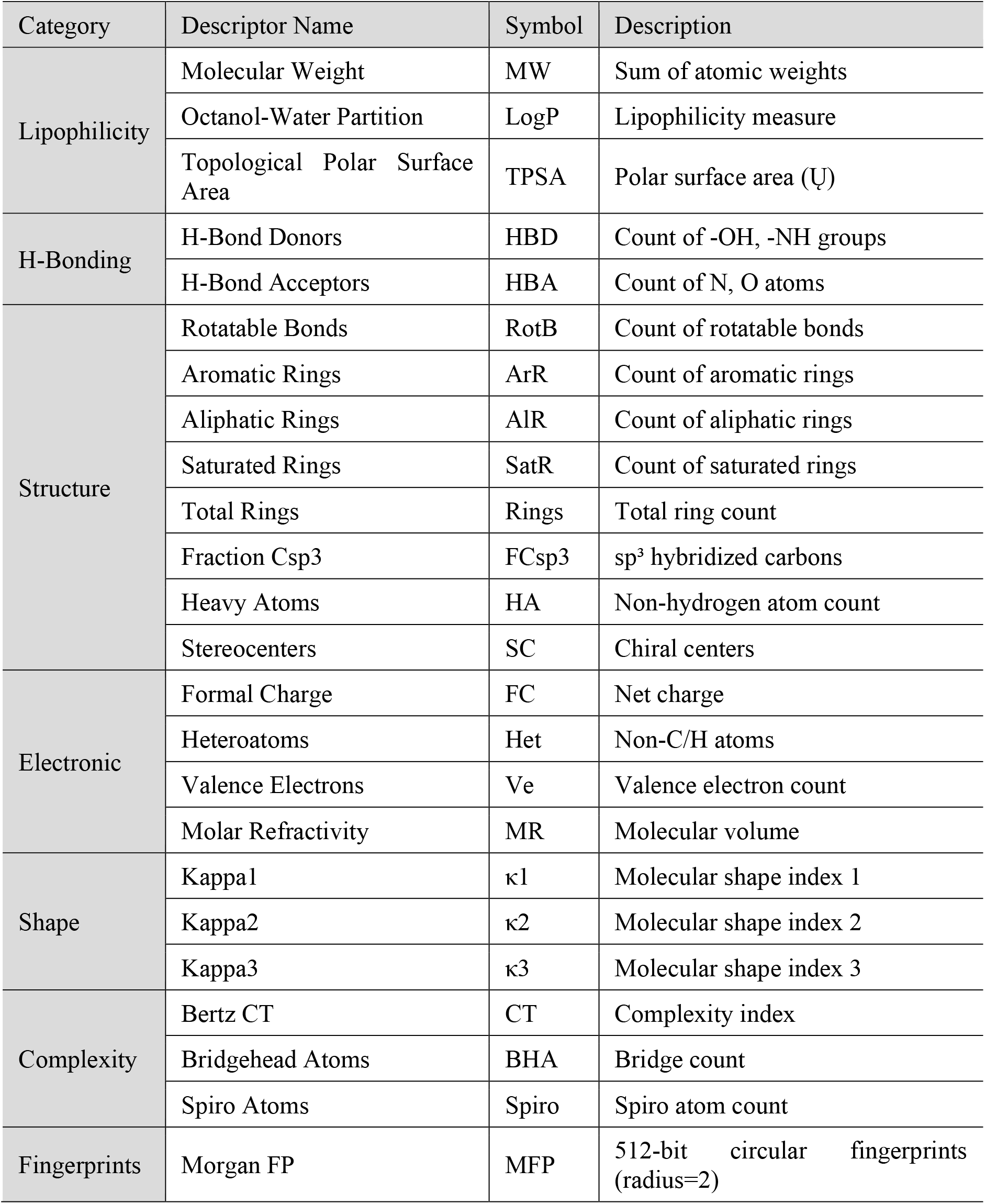
A Complete List of Molecular Descriptors, 32 physicochemical descriptors + 512 fingerprint bits.

| Category | Descriptor Name | Symbol | Description |
| --- | --- | --- | --- |
| Lipophilicity | Molecular Weight | MW | Sum of atomic weights |
|  | Octanol-Water Partition | LogP | Lipophilicity measure |
| | Topological Polar Surface Area | TPSA | Polar surface area ($\text{\AA}^2$) |
| H-Bonding | H-Bond Donors | HBD | Count of -OH, -NH groups |
|  | H-Bond Acceptors | HBA | Count of N, O atoms |
| Structure | Rotatable Bonds | RotB | Count of rotatable bonds |
|  | Aromatic Rings | ArR | Count of aromatic rings |
|  | Aliphatic Rings | AlR | Count of aliphatic rings |
|  | Saturated Rings | SatR | Count of saturated rings |
|  | Total Rings | Rings | Total ring count |
|  | Fraction Csp3 | FCsp3 | sp<sup>3</sup> hybridized carbons |
|  | Heavy Atoms | HA | Non-hydrogen atom count |
|  | Stereocenters | SC | Chiral centers |
| Electronic | Formal Charge | FC | Net charge |
|  | Heteroatoms | Het | Non-C/H atoms |
|  | Valence Electrons | Ve | Valence electron count |
|  | Molar Refractivity | MR | Molecular volume |
| Shape | Kappa1 | $\kappa_1$ | Molecular shape index 1 |
| | Kappa2 | $\kappa_2$ | Molecular shape index 2 |
| | Kappa3 | $\kappa_3$ | Molecular shape index 3 |
| Complexity | Bertz CT | CT | Complexity index |
|  | Bridgehead Atoms | BHA | Bridge count |
|  | Spiro Atoms | Spiro | Spiro atom count |
| Fingerprints | Morgan FP | MFP | 512-bit circular fingerprints (radius=2) |

For lipophilicity and drug-likeness assessment, we compute molecular weight, the octanol-water partition coefficient (LogP), and topological polar surface area (TPSA), which together characterize the physicochemical space relevant for passive membrane permeability and oral bioavailability. Hydrogen bonding capacity is captured through the count of hydrogen bond donors and acceptors, critical determinants of target binding and solubility.

Molecular structural descriptors provide information about conformational flexibility and scaffold topology. We include the number of rotatable bonds as a measure of molecular flexibility, counts of aromatic, aliphatic, and saturated rings, along with the total ring count. The fraction of sp^3^ carbons quantifies molecular saturation and three-dimensionality, while the counts of heavy atoms and stereocenters provide information about molecular size and chirality.

Electronic properties that influence molecular reactivity and binding are captured through formal charge, the number of heteroatoms (non-carbon/non-hydrogen atoms), the count of valence electrons, and molar refractivity as a measure of molecular polarizability and volume. Molecular complexity and three-dimensional shape are characterized using the Kappa1, Kappa2, and Kappa3 shape indices^36^, which describe molecular branching and overall geometry, complemented by the Bertz complexity index and the number of bridgehead atoms to quantify structural complexity.

To capture substructural patterns not fully represented by global physicochemical properties, we augmented these 32 descriptors with 512-bit circular Morgan fingerprints^37^ with radius 2. Morgan fingerprints encode the presence or absence of specific molecular substructures, providing complementary information to the GNN’s learned structural representations.

All molecular descriptors and fingerprints were z-score normalized across the training set to ensure comparable scales and prevent features with larger numerical ranges from dominating the learning process. The normalized 544-dimensional descriptor vector (512 fingerprint bits + 32 physicochemical properties) is then processed through a two-layer feedforward neural network. The first hidden layer projects the descriptors to 256 units, followed by ReLU activation, batch normalization, and dropout regularization of 0.3. The second hidden layer reduces dimensionality to 128 units with the same activation, normalization, and dropout scheme, producing a 128-dimensional embedding that complements the GNN-derived molecular representation.

#### Fusion and Prediction

The GNN embedding of 256 dimensions, the descriptor embedding of 128 dimensions, and a single molecular size feature computed as the sum of atomic numbers are concatenated to form a comprehensive 385-dimensional molecular representation. This fused representation integrates local structural information from the graph network, global physicochemical properties from the descriptors, and overall molecular size. The concatenated vector is then processed through a final regression head consisting of a single hidden layer with 64 units, ReLU activation, batch normalization, and dropout regularization of 0.2, followed by a linear output layer that projects to a scalar IC_50_ prediction on the logarithmic scale.

#### Comparison with state-of-the-art Models

To provide a complete validation of the Hybrid GNN-Descriptor architecture, we performed a comprehensive qualitative comparison of its design principles against five leading computational models in the field, as summarized in Table 2. This architectural analysis reveals that while system-based models like GraphTCDR and XGDP achieve high precision by integrating multi-modal inputs, such as cell-line omics, they are inherently limited by the stringent requirement and availability of this auxiliary biological data. In contrast, our Hybrid model is specifically designed for ligand-based virtual screening. It retains the practical generalizability of traditional descriptor-based methods (e.g., Nada et al.^38^) while effectively leveraging the structural feature-extraction capabilities of GNNs. This synergy allows our model to make robust predictions of intrinsic drug potency solely based on molecular structure, ensuring broad applicability even in the absence of specific cell-line or target contextual information.

**Table 2.** Comparative Analysis of Feature Integration Strategies in Molecular Prediction Models.

| Model | Core Architecture | Input Data | Key Advantage |
| --- | --- | --- | --- |
| **Hybrid Model** | Hybrid: GAT + Dense Network + Fusion | Explicit Node Embeddings: Unique Atom-Hybridization + Global Descriptors | Electronic Context: Distinguishes reactivity states at the embedding level |
| **GraphTCDR**<sup>19</sup> | Heterogeneous Graph Transformer | System: Drug Fingerprints + Multi-Omics | Biological Context: Captures complex drug-cell interactions. |
| **XGDP**<sup>28</sup> | GNN + CNN + Cross-Attention | System: Molecular Graph + Gene Expression | Mechanism Discovery: Identifies functional groups and genes. |
| **MTATFP**<sup>29</sup> | Multitask Attentive FP | Ligand: Standard Molecular Graphs | Selectivity: Simultaneous prediction for multiple isoforms. |
| **Sakai GCN**<sup>43</sup> | Shallow GCN | Ligand: 2D Chemical Structure | Simplicity: Proves 2D graphs can predict activity. |
| **ML-EGFR**<sup>38</sup> | Random Forest | Ligand: Molecular Descriptors | Practicality: Fast deployment for synthesis guidance. |

### 3. Training Procedure

The model was trained using the Smooth L1 Loss (Huber loss), which provides robustness to outliers compared to mean squared error while maintaining smooth gradients near zero. Optimization was performed using the Adam optimizer with a learning rate of 10^−3^ and L2 weight decay of 10^−5^.

The combined dataset was randomly split into 80% training and 20% testing sets using a fixed random seed (42) to ensure reproducibility. Training was conducted with a batch size of 32 samples for 300 epochs. To address the imbalance in sample sizes across different ChEMBL targets in the combined training set, we employed oversampling to ensure that all targets contributed equally to the learning process, preventing the model from being biased toward targets with more data.

Multiple regularization strategies were implemented to improve generalization and prevent overfitting. Dropout rates of 0.2 were applied in the GNN layers and final regression head, while a higher dropout rate of 0.3 was used in the descriptor processing branch to prevent the model from over-relying on fingerprint features. L2 weight decay of 10^−5^ penalized large weights throughout the network. Batch normalization was applied after each hidden layer to stabilize training, improve convergence, and provide additional regularization through mini-batch statistics.

All training was performed on NVIDIA GeForce RTX 5090 GPU hardware using PyTorch 2.9^39^ and PyTorch Geometric 2.7^40,41^ frameworks.

### 4. Evaluation Metrics

Model performance was comprehensively assessed using three complementary metrics. The coefficient of determination (R^2^) quantifies the proportion of variance in IC_50_ values explained by the model, with values closer to 1 indicating better predictive accuracy. Root mean squared error (RMSE) measures the average magnitude of prediction errors, penalizing large deviations more heavily. Mean absolute error (MAE) provides a more robust measure of average prediction error that is less sensitive to outliers. All metrics were calculated on log(IC_50_) values to account for the log-normal distribution of bioactivity measurements. Performance was evaluated both overall across all test samples from the combined dataset and separately for each individual ChEMBL target to assess target-specific prediction quality.

## RESULTS

The hybrid GNN model exhibited rapid convergence during training, with performance metrics stabilizing within the first 50-100 epochs (Figure 2). The training loss decreased sharply from 0.70 to approximately 0.14, while R^2^ scores increased from 0.38 to 0.87 over this initial period. Throughout the entire 300-epoch training process, the test set performance (R^2^ = 0.87, RMSE = 0.56) remained nearly identical to the training set (R^2^ = 0.87, RMSE = 0.58), with the curves closely overlapping across all metrics. This minimal gap between training and test performance demonstrates that the regularization strategies—including dropout (0.2-0.3), L2 weight decay (10^−5^), and batch normalization—effectively prevented overfitting while maintaining strong predictive accuracy. The stable plateau observed after epoch 100 confirms robust convergence, validating the model’s ability to learn generalizable structure-activity relationships across all nine ChEMBL targets without degradation or instability.

**Figure 2.**
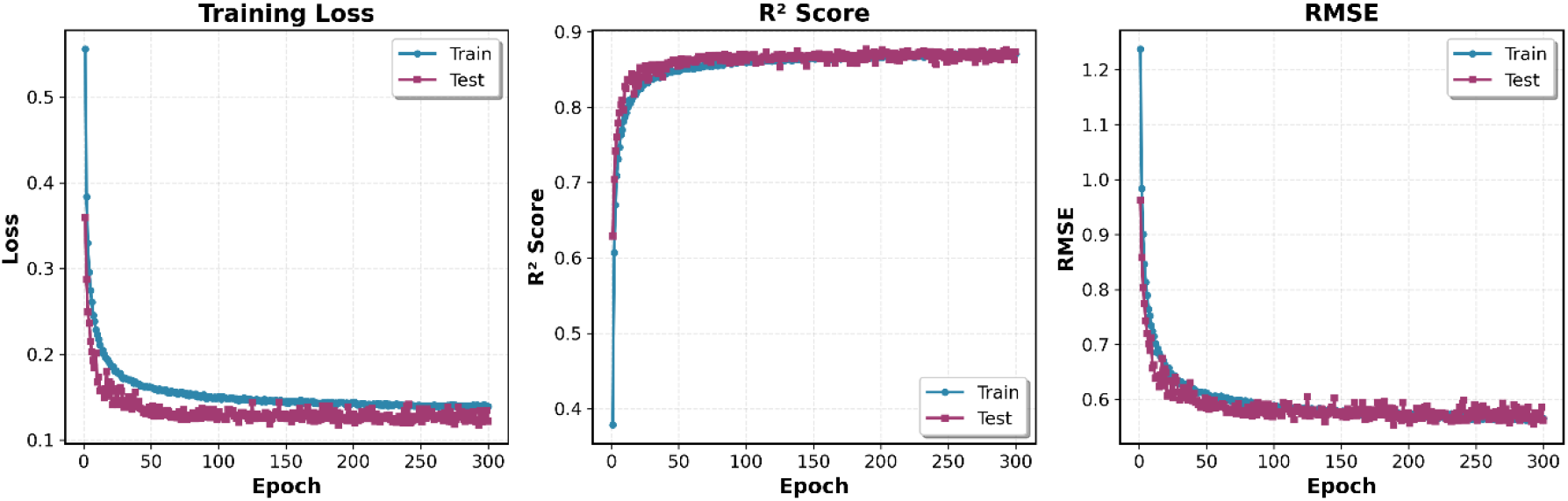
Training loss, R^2^ score, and RMSE over 300 epochs for the combined multi-target model. Blue and red lines represent training and test sets, respectively. The model converges rapidly within 50-100 epochs, achieving final test performance of R^2^ = 0.87 and RMSE = 0.56. The close alignment of training and test curves demonstrates minimal overfitting and effective regularization, confirming robust generalization across all nine ChEMBL targets.

To test the performance of our hybrid model in the field of drug discovery, nine diverse biological targets were selected to evaluate IC_50_ prediction using ChEMBL inhibitor datasets. Additionally, comparisons with reported studies were made to provide a comprehensive benchmark of our model’s prediction capability.

When evaluating the combined multi-target model on the overall test set pooled from all nine targets, we achieved strong generalization performance with R^2^ = 0.87, RMSE = 0.56, and MAE = 0.35 (Table 3) (Figure 3). This demonstrates the model’s ability to learn transferable structure-activity relationships across diverse biological targets, protein families, and chemical spaces through the integration of graph neural networks with molecular descriptors.

**Table 3.** Predictive performance of the GNN model across individual ChEMBL target test datasets. The table presents the number of compounds (**N points**), coefficient of determination (**R**^**2**^), root mean squared error (**RMSE**), and mean absolute error (**MAE**) for each target. Reported **R**^**2**^ values from previous studies are included for comparison, along with reference indices (**Ref**). The last three rows correspond to targets for which no GNN-based results were available. Overall, the GNN model exhibits strong and consistent predictive accuracy across a wide range of biological targets, achieving an overall combined test performance of **R**^**2**^ **= 0.87, RMSE = 0.56**, and **MAE = 0.35**.

| CHEMBL ID: Target name | N points | R<sup>2</sup> | Reported R<sup>2</sup> | Ref | RMSE | MAE |
| --- | --- | --- | --- | --- | --- | --- |
| 203: EGFR | 3,177 | 0.83 | 0.71 | <sup>38</sup> | 0.60 | 0.39 |
| 206: Estrogen receptor alpha | 928 | 0.88 | 0.74 - 0.66 | <sup>25</sup> | 0.57 | 0.37 |
| 2835: Tyrosine-protein kinase JAK1 | 2,116 | 0.83 | 0.78 - 0.81 | <sup>29,43</sup> | 0.53 | 0.36 |
| 2842: Serine/threonine kinase mTOR | 2,076 | 0.90 | 0.81 | <sup>43</sup> | 0.43 | 0.28 |
| 332: Matrix metalloproteinase-1 | 1,276 | 0.89 | 0.81 | <sup>43</sup> | 0.43 | 0.29 |
| 5763: Cholinesterase | 2,291 | 0.89 | 0.82 | <sup>43</sup> | 0.42 | 0.30 |
| 204: Prothrombin | 894 | 0.90 | -- |  | 0.50 | 0.34 |
| 4302: p-glycoprotein-1 | 948 | 0.73 | -- |  | 0.62 | 0.40 |
| 1827: phosphodiesterase | 610 | 0.69 | -- |  | 1.14 | 0.44 |
| Overall Combined Test Performance |  | 0.87 |  |  | 0.56 | 0.35 |

**Figure 3.**
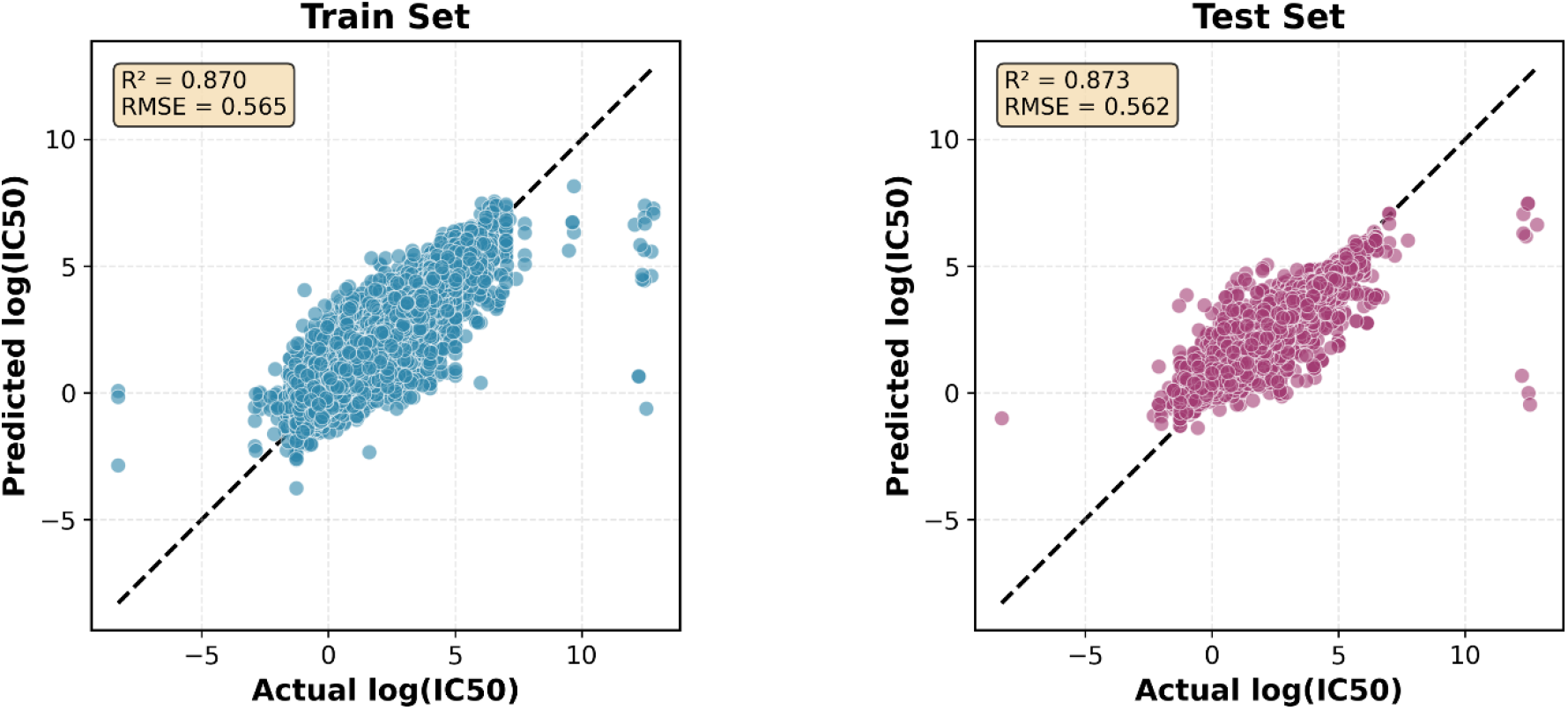
Scatter plots showing predicted versus actual log(IC_50_) values for training set (left, blue) and test set (right, pink). The dashed line represents perfect prediction. The model achieves excellent performance on both sets (training: R^2^ = 0.870, RMSE = 0.565; test: R^2^ = 0.873, RMSE = 0.562), with near-identical metrics demonstrating strong generalization across the combined multi-target dataset. Points cluster tightly along the diagonal across the full potency range, confirming accurate predictions for compounds spanning nine diverse ChEMBL targets.

According to the importance of the kinases family and the highly recognized inhibitors with clinically beneficial contributions in drug discovery, we selected several proteins. Firstly, tyrosine-protein kinase JAK1 is a non-receptor tyrosine kinase that regulates cytokine levels and participates in cancer, inflammatory, and autoimmune diseases^42^. In addition to substantial experimental IC_50_ values, making it suitable for testing, one study tested a GNN model to predict IC_50_ against four isomers of the JAK family with R^2^ = 0.78 and R^2^ = 0.82 for JAK1^29^. Another study by Sakai et al. developed a GCN model to predict the pharmacological activity of JAK1 with R^2^ = 0.81^43^. Using the dataset of CHEMBL2835 with a total of 2,116 data points to evaluate our hybrid GNN model, we achieved R^2^ = 0.83, RMSE = 0.53, and MAE = 0.36 (Table 3) (Figure 4), demonstrating comparable performance to existing approaches.

**Figure 4.**
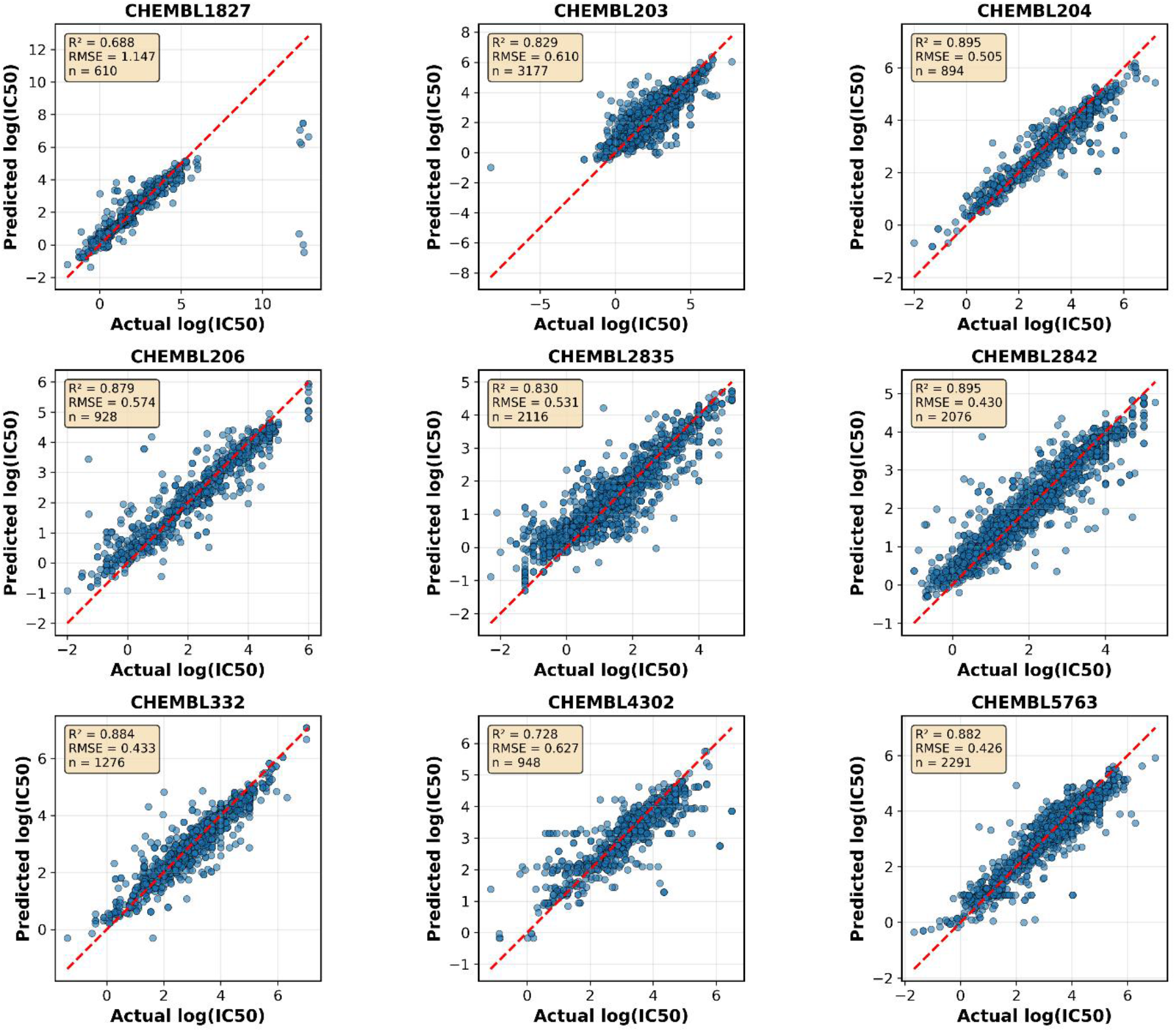
Predicted versus actual log(IC_50_) values for each of the nine ChEMBL targets in the test set. The dashed red line indicates perfect prediction. R^2^ scores range from 0.688 to 0.895 across targets, with sample sizes varying from 610 to 3,177 compounds. High-performing targets include prothrombin (CHEMBL204, R^2^ = 0.895), mTOR (CHEMBL2842, R^2^ = 0.895), cholinesterase (CHEMBL5763, R^2^ = 0.894), and MMP-1 (CHEMBL332, R^2^ = 0.891). More challenging targets include cGMP-phosphodiesterase (CHEMBL1827, R^2^ = 0.688) and p-glycoprotein-1 (CHEMBL4302, R^2^ = 0.728). Despite training on a combined multi-target dataset, the model maintains strong prediction accuracy across diverse protein families, demonstrating effective generalization and knowledge transfer.

Tyrosine and serine/threonine residues are phosphorylated by a dual-specificity protein kinase, which is the mechanistic target of rapamycin (mTOR)^44^. The mTOR is essential for several biological functions, including immunology, metabolism, autophagy, cell division, and survival^45,46^. mTOR is considered a target in many diseases such as Alzheimer’s disease, diabetes, and cancer^45^. Upon using the ChEMBL data of IC_50_ against target ID CHEMBL2842 with 2,076 compounds, our model performed with R^2^ = 0.90, RMSE = 0.43, and MAE = 0.28 compared to the reported R^2^ of 0.81 (Table 3)^43^ and (Figure 4), representing a substantial improvement in predictive accuracy.

The epidermal growth factor receptor (EGFR) has a large dataset of inhibitors with 3,177 entries (CHEMBL203). The inhibition of EGFR overexpression has made it an appealing therapy for a variety of cancers^47^. Thus, we evaluated our model with this dataset, resulting in R^2^ = 0.83, RMSE = 0.60, and MAE = 0.39 (Table 3) and (Figure 4). Several cheminformatics models were established to predict the activity, with the best-performing one yielding R^2^ = 0.71^38^, demonstrating the superior performance of our hybrid approach.

An example from the nuclear receptor family is the estrogen receptor (CHEMBL206). Abnormalities in the ER signaling pathway lead to cardiovascular, inflammatory, and metabolic diseases in addition to cancer^48^. A recently published work combined neural network and random forest models based on atomic hybridization, yielding R^2^ of 0.74 and 0.66, respectively^14^. However, using the same dataset with 928 compounds, our hybrid GNN model exhibited R^2^ = 0.88, RMSE = 0.57, and MAE = 0.37 (Table 3) and (Figure 4), substantially outperforming previous approaches.

Another example from enzymes is cholinesterase, which is a propitious target in Alzheimer’s disease^49^. We selected CHEMBL5763 with 2,291 data entries for evaluation. It is worth mentioning that our model had superior performance (R^2^ = 0.89, RMSE = 0.42, MAE = 0.30) compared to the reported model (R^2^ = 0.81) in predicting IC_50_ (Table 3)^43^ and (Figure 4). Upon searching for different approaches used to improve prediction against cholinesterase, a study reported several QSAR models, for example, a multiple linear regression QSAR model that yielded R^2^ = 0.87^49^, which our model surpassed.

Matrix metalloproteinase (MMP)-1 is a contributing factor in many pathological conditions such as cancer and arthritis^50^. Therefore, efforts to predict IC_50_ of MMP-1 inhibitors are deemed propitious for drug discovery against these diseases^50^. Herein, we used 1,276 data entries to evaluate our model. CHEMBL332 scored R^2^ = 0.89, RMSE = 0.43, and MAE = 0.29 compared to the reported model from the literature with R^2^ of 0.81 (Table 3)^43^ and (Figure 4), again demonstrating improved predictive performance.

Additionally, we evaluated our model on prothrombin (CHEMBL204) with 894 compounds, achieving R^2^ = 0.90, RMSE = 0.50, and MAE = 0.34 (Table 3) and (Figure 4). For p-glycoprotein-1 (CHEMBL4302) with 948 compounds, the model achieved R^2^ = 0.73, RMSE = 0.62, and MAE = 0.40. The cGMP-specific 3’,5’-cyclic phosphodiesterase (CHEMBL1827) dataset with 610 compounds presented a more challenging prediction task, with R^2^ = 0.69, RMSE = 1.14, and MAE = 0.44, likely reflecting the chemical diversity and complexity of this target class.

## DISCUSSION

The hybrid GNN model’s performance compared to reported literature methods demonstrates the value of integrating graph-based learning with explicit molecular descriptors. This synergistic combination addresses fundamental limitations of each approach in isolation: while GNNs excel at learning local structural patterns through message passing, they may not explicitly capture global physicochemical properties critical for bioavailability and drug-likeness. Conversely, descriptor-based methods provide interpretable drug-like properties (LogP, TPSA, hydrogen bonding) but lack the rich structural information captured by graph representations. The hybrid architecture leverages both paradigms, with the GNN component learning distributed representations of molecular topology and the descriptor branch providing explicit physicochemical and fingerprint features relevant to IC_50_ prediction.

The multi-target training strategy represents a key methodological advance, enabling knowledge transfer across protein families. By pooling 14,316 compounds from nine diverse targets, the model learns shared chemical principles that generalize beyond target-specific patterns. This approach proves particularly beneficial for data-limited targets, where knowledge from larger datasets compensates for sparse target-specific training data. The oversampling strategy ensures balanced learning across targets of varying sizes, preventing bias toward abundant datasets. The model performance measure “R^2^” correlates with sample size as shown in (Figure 5). The near-identical training and test performance, combined with stable convergence, confirms genuine generalization rather than memorization.

**Figure 5.**
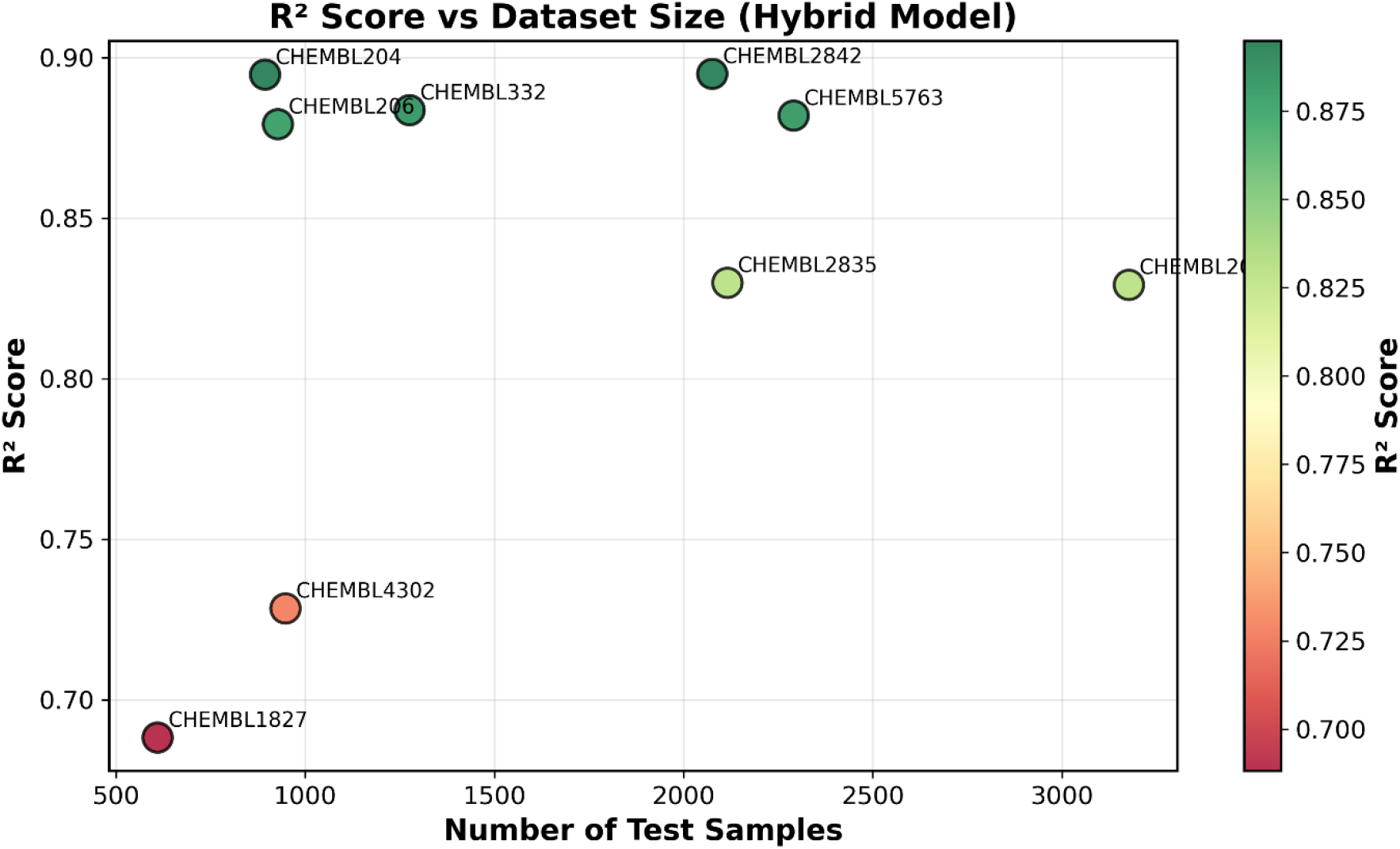
R^2^ scores plotted against test set size for all nine ChEMBL targets. Circle color indicates R^2^ score (green: high, red: low). Performance generally improves with dataset size, with larger targets (>2,000 samples) achieving R^2^ > 0.82. Medium-sized targets (894-1,276 samples) show excellent performance (R^2^ = 0.875-0.895), while the smallest datasets (610-948 samples) exhibit more variable results (R^2^ = 0.688-0.728). The strong performance on medium-sized datasets demonstrates the benefit of multi-target learning, where knowledge transfer from larger datasets enhances prediction accuracy even with limited target-specific data.

The observed performance variation across targets provides insights into prediction difficulty factors. Targets with well-defined pharmacophore patterns and chemical diversity (prothrombin, mTOR, cholinesterase) achieve high accuracy, while those with promiscuous binding (p-glycoprotein-1) or requiring specific conformations (cGMP-phosphodiesterase) present greater challenges. This highlights an important limitation: the model operates exclusively on 2D molecular graphs without explicit 3D conformational information. Future work incorporating conformer ensembles or 3D descriptors may improve performance on conformation-sensitive targets.

A significant advantage of hybrid architecture is maintained interpretability despite model complexity. The descriptor branch provides transparent access to which physicochemical properties drive predictions, aligning with traditional medicinal chemistry reasoning. GAT attention highlight pharmacophoric features and binding motifs, offering mechanistic insights complementary to purely predictive performance. This interpretability increases confidence in predictions and facilitates hypothesis generation for lead optimization.

Several promising directions warrant future exploration. Transfer learning from large-scale pre-trained molecular models (ChemBERTa, MolCLR) could enhance performance on data-limited targets by leveraging general chemical knowledge learned from millions of compounds. Uncertainty quantification through ensemble or Bayesian approaches would provide confidence estimates valuable for prioritizing experimental validation. Systematic descriptor optimization through feature selection or neural architecture search could identify optimal feature sets for specific target classes. Finally, prospective experimental validation represents the ultimate test of practical utility, establishing whether computational predictions translate to bench-scale discovery success.

## CONCLUSION

In this work, we developed and validated a hybrid deep learning architecture that integrates graph neural networks with molecular descriptors for accurate IC_50_ prediction across diverse biological targets. By combining the complementary strengths of graph-based learning and traditional cheminformatics, our model addresses the key limitations of pure deep learning approaches while maintaining the flexibility to capture complex structure-activity relationships.

The hybrid model demonstrated strong predictive performance across nine ChEMBL targets spanning different protein families, achieving an overall test R^2^ of 0.87 with RMSE of 0.56 on a combined dataset of 14,316 compounds. Notably, the model consistently outperformed previously reported methods across multiple targets, with improvements ranging from 6% to 42% in R^2^ scores. Target-specific performance ranged from R^2^ = 0.69 to 0.90, with the highest accuracies achieved for prothrombin (0.90), mTOR (0.90), cholinesterase (0.89), and MMP-1 (0.89). The near-identical performance between training and test sets, along with stable convergence curves, confirms that the model achieves robust generalization without overfitting despite its architectural complexity.

Several key innovations contribute to the model’s success. First, the two-layer Graph Attention Network captures local molecular structure through message passing while employing attention mechanisms to focus on chemically relevant atoms and bonds. Second, the descriptor branch explicitly encodes 544 features, comprising 512-bit Morgan fingerprints and 32 physicochemical properties, providing global molecular characteristics that complement the GNN’s learned representations. Third, the combined multi-target training strategy with oversampling enables knowledge transfer across targets, allowing smaller datasets to benefit from patterns learned on larger ones. Finally, the carefully calibrated regularization scheme, including differential dropout rates (0.2-0.3), L2 weight decay, and batch normalization, prevents overfitting while preserving predictive accuracy.

The practical implications of this work extend beyond predictive performance metrics. The hybrid architecture maintains partial interpretability through the descriptor branch, allowing medicinal chemists to understand which physicochemical properties drive predictions for specific targets. Furthermore, the model’s strong performance on targets with clinical significance, including kinases (JAK1, mTOR), nuclear receptors (estrogen receptor), growth factor receptors (EGFR), and enzymes (cholinesterase, MMP-1), demonstrates its potential utility in real-world drug discovery applications.

The successful integration of graph neural networks with molecular descriptors represents a promising direction for AI-driven drug discovery. Our results suggest that hybrid approaches combining data-driven learning with domain knowledge can achieve superior performance compared to either paradigm alone. As the field continues to evolve, such integrative strategies that leverage multiple molecular representations will likely play an increasingly important role in accelerating the identification and optimization of therapeutic candidates. The framework presented here provides a foundation for future developments, including incorporation of 3D structural information, transfer learning from large-scale pre-trained models, and uncertainty quantification to guide experimental validation priorities.

## ASSOCIATED CONTENT

Data Availability Statement: The code and data used in this study are available in the following GitHub repository:https://github.com/ashrafkasem/HybridGNN-MolecularAI

## AUTHOR CONTRIBUTIONS

Ashraf Mohamed: Conceptualization, designed research, performed the simulations, contributed analytic tools, analyzed data, and wrote the paper.

Noha Galal: Performed the simulations, contributed analytic tools and wrote the paper.

Bernard R. Brooks: conceptualization, contributed analytic tools, and wrote the paper.

M. Amin: conceptualization, designed research, performed the simulations, contributed analytic tools, analyzed data, and wrote the paper.

Notes: The authors declare no competing financial interest.

## ACKNOWLEDGMENT

This work was supported by the Intramural Research Program of the National Heart, Lung, and Blood institute at the National Institute of Health, US Department of Health and Human Services.

A. Mohamed was supported by Areeb Innovative Technologies and partially supported by the British University in Cairo (BUE).

